# Higher rewards lead to more accurate flower detection and increased contrast sensitivity in the bumblebee *Bombus terrestris*

**DOI:** 10.64898/2026.08.26.747241

**Authors:** Théo Robert, Emily Flett, Hugo Le Lay, Marin Nicolas, Vivek Nityananda

## Abstract

In vertebrates, top-down visual attention is a cognitive process where internal goals modulate the tuning of peripheral sensory systems. This leads to increased perceived contrast to both goal-relevant objects and areas of the visual field that are attended. Such a system would also be beneficial to bees, enabling them to detect and recognise the most profitable flowers in their environment. We tested whether bumblebees possess a top-down attentional system resembling that seen in vertebrates. We trained two groups of bees to collect rewards under high contrast targets. To potentially induce a difference in attention while searching for the targets, one group received a higher concentration of sucrose rewards compared to the other. During tests, the targets were presented with a series of lower contrasts to measure the contrast sensitivity curves of the bees induced by the different learnt reward levels. We predicted a stronger effect of any attention-like process on contrast sensitivity in the high reward group. We also repeated this experiment with the neonicotinoid pesticide imidacloprid dissolved in the sucrose rewards to test whether this affects bee attention. Across all test contrasts, higher rewards significantly increased bee accuracy when locating targets, lowered contrast thresholds and reduced the latency to make first choices. Imidacloprid reduced bee accuracy but did not influence first choice latency. These results suggest that learnt floral rewards can influence bee behavioural contrast sensitivity in a manner resembling vertebrate top-down attention and that imidacloprid may modulate this through effects on their nervous system.

## Introduction

Animals are constantly exposed to multiple simultaneous sensory stimuli, and it is impossible for them to process all the sensory information they receive. To tackle this problem, they have evolved cognitive processes that filter and prioritise important information from noise. These processes are generally described as attention, which can involve different mechanisms (for reviews, see Carrasco, 2011; Petersen and Posner, 2012). Two key mechanisms have been categorised as bottom-up and top-down attention. In bottom-up or exogenous attention, attention is automatically captured by sudden or salient cues in the environment (Chica et al., 2013). In top-down or endogenous attention, attention is directed toward an area in space as a result of learnt goals (Chica et al., 2013).

Top-down visual attention was shown in a classic experiment (Posner, 1980) by presenting participants an arrow as a cue pointing to one side of a screen to indicate where a target would appear. The presence of the cue led to them detecting a target faster if it appeared on the side of the screen indicated by the cue than on the other side. Top-down attention can also be deployed when individuals look for a specific object in their environment in what is called feature-based attention. Here, the sensory system of the subject is tuned to improve the detection of specific features previously associated with the object such as shape, colour, orientation or movement direction (Herrmann et al., 2012; Ling et al., 2009; Liu et al., 2007; White and Carrasco, 2011; Zhou and Desimone, 2011; for a reviews, see Liu, 2019; Scolari et al., 2014).

An important characteristic of top-down attention in humans is the fact that it can transiently increase the contrast sensitivity of the subject in the region towards which the attention was directed (Barbot et al., 2012; Herrmann et al., 2012; Lee and Carrasco, 2025; Ling and Carrasco, 2006; Liu et al., 2009; Pestilli et al., 2011). For example, displaying a bar pointing to where a subsequent target will briefly appear increases contrast sensitivity in the corresponding area of the visual field, allowing individuals to detect the target at a lower contrast than without the cue (Ling and Carrasco, 2006). Similarly, indicating the correct orientation of subsequent targets allows individuals to use their feature based attention to better perceive them at low contrasts (Herrmann et al., 2012).

Numerous studies have demonstrated top-down attention in both human and non-human primates (e.g. Busse et al., 2008; Chica et al., 2013; Di Bello et al., 2025; Fernández et al., 2022; Jigo and Carrasco, 2020; Mahadevan et al., 2018; Pham et al., 2018). It also seems to be present in other mammals such as rats (Marote and Xavier, 2011; Rosner and Mittleman, 1996), and in birds such as chickens (Sridharan et al., 2014) and crows (Quest et al., 2022), suggesting it might have evolved convergently in multiple animals. An ability to improve target detection based on previously learned information would also be a very useful mechanism for bees. These insects forage on flowers that vary in colour and shape, providing clear cues to reward, while floral reward value influences motivation and learning. Top-down attention based on learnt cues could therefore help bees locate and detect flowers based on their reward value.

To test this, we therefore asked whether bumblebees have a cognitive system resembling top-down attention in vertebrates. We trained two groups of bumblebees to detect and collect a sucrose solution reward under 6 green targets on a computer screen. The reward for one group of bees had a higher concentration than for the other group, potentially creating a difference in any learnt top-down mechanisms. In subsequent tests, the targets were presented with lower contrasts than during training and we measured the bees’ accuracy and latency to choose the target locations. Higher reward concentration leads to greater foraging motivation in bees (Dornhaus and Chittka, 2005; Frasnelli et al., 2021; Lihoreau et al., 2011; Muth et al., 2015; Pattrick et al., 2023; Whitney et al., 2008). We therefore hypothesized that the bees in the higher reward concentration group would be more motivated, leading to higher top-down attention towards the targets, in turn allowing the bees to detect them at a lower contrast than the other group of bees.

To further understand ecological and physiological effects on this process, we investigated how exposure to the neonicotinoid pesticide imidacloprid affected any top-down mechanisms observed.

Exposure to sublethal doses of neonicotinoid pesticides has been shown to hinder the foraging abilities of several species of bees through the disturbance of numerous cognitive functions. For example, these pesticides can affect bees’ learning and memory (Ke et al., 2023; Mustard et al., 2020; Muth et al., 2019; Stanley et al., 2015; Tison et al., 2019; Wright et al., 2015), but also their navigation (Fischer et al., 2014; Samuelson et al., 2016; Tison et al., 2016) or their motivation to forage (Lämsä et al., 2018; Muth and Leonard, 2019). We therefore asked whether imidacloprid, a nicotinoid, could hinder the attentional processes of bumblebees leading to a degradation of their ability to detect the targets. In addition, impidacloprid targets acetylcholine receptors in insects (Brown et al., 2006; Liu et al., 2005; Matsuda et al., 2020; Taillebois et al., 2018) and we could therefore use it to test for any potential role of cholinergic receptors in top-down attention in insects. To test this, we repeated our protocol with a sublethal dose of imidacloprid dissolved in the sugar rewards throughout the experiment.

## Material and Methods

### Animals and Setup

In our experiment, we used bumblebee workers (*Bombus terrestis*) from 6 commercially reared colonies obtained from Agralan Ltd, UK. Upon their arrival at the laboratory at Newcastle University, the colonies were transferred to a plastic nest box (L=28 cm, W=16 cm, H=12 cm; Fig. 1A) divided in two chambers (L=16 cm, W=14 cm, H=12 cm). The brood was placed in one chamber while the second chamber contained cat litter where the bees could deposit their waste. This last chamber was connected to a transparent square shaped acrylic tunnel (L=60 cm, S=5 cm) leading to a foraging arena (L=45 cm, W=60 cm, H=40 cm). The tunnel had several plastic doors allowing the experimenter to select and release bees one by one in the arena. The floor of the arena was covered with a red and white random checkerboard pattern which provided optic flow to help the bees stabilise their flight (Linander et al., 2017; Linander et al., 2018). The wall of the arena opposite to the tunnel entrance consisted of a computer screen (Dell S2419HGF, LCD, 1080p, 144 Hz) allowing us to display stimuli during the experiment. This screen was chosen as it was flicker free with a refresh rate faster than the bees’ flicker fusion (Meyer-Rochow, 1981; Skorupski and Chittka, 2010; Srinivasan and Lehrer, 1984; Srinivasan and Lehrer, 1985). The arena was covered with an acrylic lid transparent to white and UV light. A full daylight spectrum light tube (Philips, Master TL5 HE 35W, 6500K) fitted to a high frequency lighting system (Philips, HF-P 1 14-35 TL5 HE III, >42KHz) was suspended over the arena. A smartphone (Xiaomi Redmi Note 11 Pro, China) was also placed horizontally over the arena during the experiment to record the test trials.

When the experiment was not ongoing, during evenings and weekends, the colony could access the arena freely. We placed gravity feeders filled with 20% w/w sucrose solution in the arena, so the bees had food ad libitum. At all times, the colony had access to pollen in a little plate placed on top of the cat litter in the nest chamber.

### Experimental Protocol

Before starting the experiment, the arena was emptied of all bees and the feeders were removed. We placed three transparent acrylic shelves against the screen (Fig. 1), equally spaced vertically with a distance of 10 cm between each shelf. These were used to support transparent plastic squared chips (L=2.5 cm, H=0.5 cm, Fig. 1) with a well at their centre that could be filled with some liquid to give sucrose solution rewards to the bees. Four chips were placed on each shelf with a distance of 14 cm between them (Fig. 1C)

Individual worker bees were collected at the end of the tunnel with a plunger and a coloured and numbered tag was glued to their thorax using superglue. Each individual was then assigned to one of two groups. In the “high motivation group” bees received 50% (w/w) sucrose as high reward for a correct choice and 20% (w/w) sucrose as low reward for a wrong choice. In the “low motivation group” bees received 30% (w/w) sucrose as a high reward for a correct choice and 20% (w/w) as low reward for a wrong one. Bees have been suggested to equally value flowers based on their ranked comparative reward value to other flowers in their environment, such that flowers with greater reward than others are equally valued regardless of their actual concentration (Solvi et al., 2022). We therefore provided 20% sucrose rather than water in the non-target flowers as a baseline to create a greater contrast (50% vs 20% sucrose) for the high motivation group than for the low motivation group (30% vs 20% sucrose). This ensured that regardless of how bees valued target flower rewards (through absolute concentration or relative to the 20% sucrose flowers) the high motivation group would value higher rewards more than the low motivation group. To avoid any bias, the experimenters running the study only knew they were using a high and low reward solution and did not know the actual concentration of the solution being used (50% or 30%) for the high rewards.

### Pretraining

In the first pretraining trial, the bee was placed on an additional transparent chip placed at the centre of the middle shelf, in front of a black screen and filled with 20 µL of 20% sucrose solution. This served to teach the bee that these chips contain rewards. After the bee had drunk the sucrose, she was allowed to take off and fly in the arena. If she did not return to her nest on her own, she was captured and returned to the colony. The chip was replaced by a new one filled with 20 µL of the high reward corresponding to the bee’s group to prepare for a second pretraining trial. During this trial, a target was displayed on the screen above the chip. This target consisted of a full contrast green (RGB values passed to the screen: 0, 1, 0 ; irradiance = 0.2 µW/cm^2^/nm) circle of 5.5 cm diameter. Bees have been shown to use achromatic cues to detect relatively small targets and to perform achromatic tasks using their green channel (Giurfa and Vorobyev, 1997; Giurfa et al., 1996; Hempel De Ibarra et al., 2001). Therefore, the use of green targets ensured that the bees would learn, identify and perceive the changes in target contrast achromatically. Once the bee returned to the tunnel from the nest, it was captured again and placed on the new chip. The rest of the second pretraining trial was similar to the first one.

### Training

After a bee experienced two pretraining trials, we began the training phase. For this phase, we placed four equally spaced chips on each of the three shelves, creating twelve possible choice locations. Above six of these locations chosen randomly, we displayed a full contrast green circle target (irradiance = 0.2 µW/cm^2^/nm), similar to the one used in the second pretraining on the computer screen. Above the six remaining locations, the screen remained black. The chips below the targets were filled with 20 µL of the high reward sucrose solution corresponding to their motivation group (50% sucrose for the high motivation group and 30% for the low motivation group). This volume was chosen to ensure that most bees would need to visit six chips before being full and returning to their nest to empty their crop. The chips that were not under targets were filled with the same amount of 20% sucrose solution. The bee was then released in the arena, and we noted the choices she made. A choice was counted when the bee landed on a chip and probed its well with her antennae or her proboscis. The training trial ended when the bee made at least six choices and returned to the colony. A bee was allowed to proceed to the test phase once she had participated in a minimum of four training bouts and had 80% correct choices on her last 20 choices.

### Testing

The test setup was the same as during training, but all chips were filled up with 20 µL of distilled water. During a single test trial, all six targets on the screen were presented at one of nine irradiance levels (a proxy for target contrast), with increasing irradiances corresponding to increased contrast against the black background. We measured the irradiances with a spectrometer (FLAME-S-UV-VIS-ES, Ocean Optics, USA). The nine irradiances used were: 0.01, 0.02, 0.03, 0.04, 0.06, 0.08, 0.10, 0.12, 0.16 µW/cm^2^/nm. Each irradiance was used for all the targets of one test trial, and the trials were presented to each bee in a random order. The bee had to make at least six choices to complete a test trial. Because the bee did not receive any reward, she would not fill her crop and thus would often not return to her nest on her own. She would then need to be captured and returned to her colony.

Between test trials, we conducted refresher trials, identical to the training ones to keep the bee motivated. A refresher was considered successful if the bee completed at least six choices and achieved at least 80% correct choices. If these conditions were not met, the refresher was repeated. All test trials were recorded at 30 frames per second.

After every trial, the chips were cleaned with 70% ethanol to remove any pheromone marking and rinsed with distilled water to remove the alcohol.

In total, 58 bees started the training process (24 in the high motivation group and 34 in the low motivation group), and 40 completed it (20 in each motivation group) and started the test phase.38 bees could complete the nine irradiance test conditions (20 in the high motivation group and 18 in the low motivation group).

**Figure 1:**
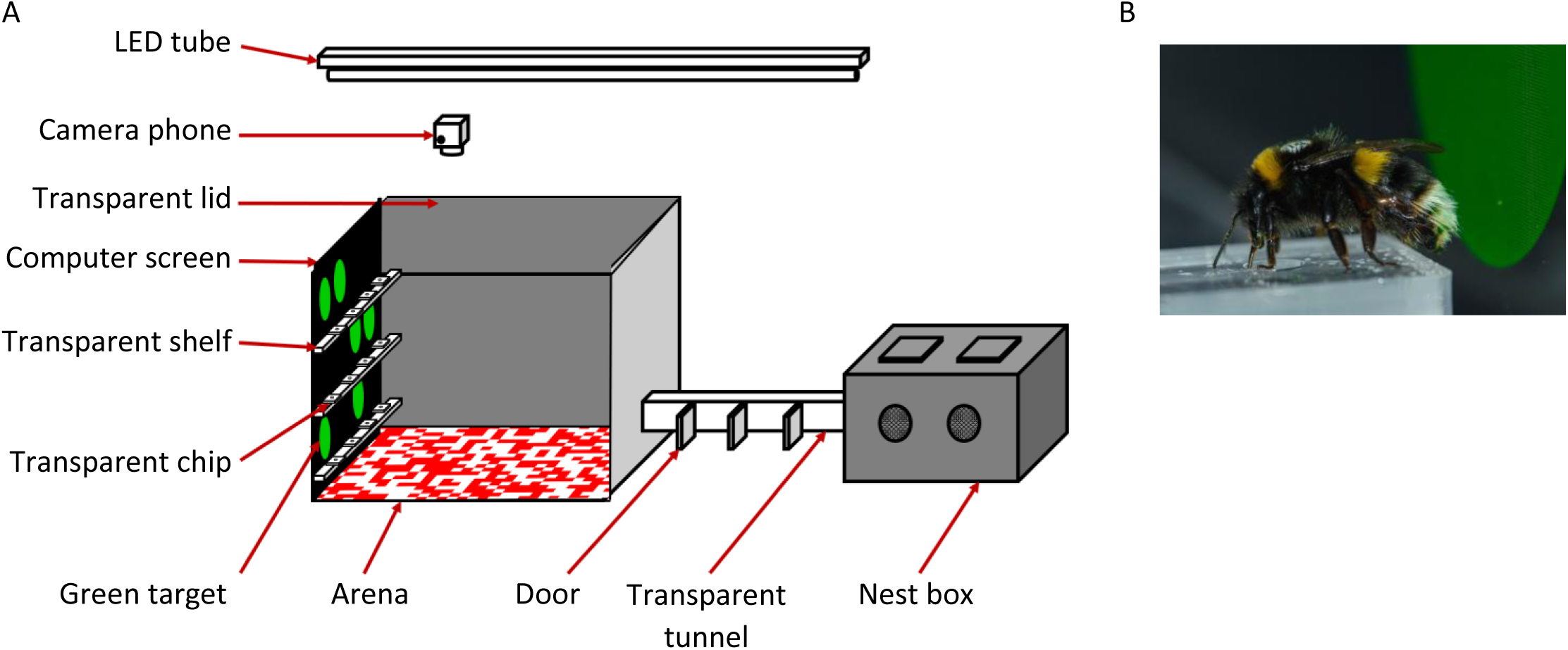
Schematic diagram of the experimental setup. A) Nest box and arena. The bumblebee colony was contained in the nest box and bees could access the experimental arena through the transparent tunnel. The experimenter could select which bee passed through using the multiple doors. A red and white pattern was displayed on the arena floor to provide optic flow to the bees. Targets could be displayed in twelve locations on the computer screen placed at the end of the arena. Three transparent shelves were placed in front of the screen, each supporting four transparent chips marking the twelve possible target locations. The chips had a little well at their centre in which we could place sugar rewards. B) Photo of a tagged bumblebee collecting a sugar reward in the well of a chip in front of a target (credit: Hugo LeLay).

### Pesticide experiment

The same protocol as above was repeated with a sublethal dose of imidacloprid, a neonicotinoid dissolved in the sucrose solutions to examine how this pesticide would affect bee attention to the targets. We added the pesticide at a concentration of 10 µg/L in all sucrose solutions. This dose was comparable to the level of pesticides found in flowers and nectar in the field (Bonmatin et al., 2003; Bonmatin et al., 2005; Byrne et al., 2014; Krischik et al., 2007; Stoner and Eitzer, 2012). We included the pesticide in both rewarding and non-rewarding solutions to make the experiment more ecologically relevant, simulating different flowers that contained pesticide. Bees in this experiment were thus exposed to the pesticide during their pretraining, training and the refresher bouts between tests.

For this experiment, 23 bees started training (8 in the high motivation group and 15 in the low motivation group) and 19 completed it and moved to the test phase (all 8 high motivation bees and 11 low motivation bees). 16 bees completed all nine test conditions (all 8 high motivation bees and 8 bees in the low motivation group).

### Video and statistical analyses

We analysed the first six choices of each bee in each test trial, because there were six possible correct locations marked by the six targets. We then analysed the proportion of these choices in which the bees probed a chip under a target.

The video recordings of the test trials were analysed to extract the bees’ latency to make their first choice. To do so, we measured the time between the video frame when the bee was no longer touching the floor of the arena upon taking off and the frame when she touched the chip that was her first choice. We focussed on first choice latency as this is the choice the bees had to make when furthest away from the screen. Thus, this was the choice when the perceived contrast of the targets may have had its strongest effect. We also similarly analysed the proportion of first choices in which the bees chose a chip below a target.

Statistical analyses were conducted with R (version 4.6.1, R Core Team, 2026) using Generalised Linear Mixed Models fitted with the package glmmTMB (Brooks et al., 2017). To improve the fit of all the models, we log transformed the target irradiance variable. To analyse the proportion of correct choices (the first choice or the first six choices), we used GLMMs with a binomial family and a logit link function. We included the target irradiance (as a continuous variable), the motivation group (two levels: high motivation and low motivation), the pesticide group (two levels: pesticide and no-pesticide) as independent variables and interaction effects between all three variables. The models also included bee identity as a random effect. The model and hypothesis were preregistered at https://aspredicted.org/jx9e9c.pdf but without the inclusion of the pesticide as a variable.

The model fitted on the probability of correct first six choices was used to generate the psychophysics curves of the bees’ contrast sensitivity (Fig. 3 and Fig. 4). To do so, we plotted the curves predicted by our model for the four conditions (High Motivation/No Pesticide, High Motivation/Pesticide, Low Motivation/No Pesticide, Low Motivation/Pesticide). Following the guidelines from Balestrucci and colleagues (Balestrucci et al., 2022), we used the function pseMer() from their MixedPsy R package to calculate the bees’ contrast sensitivity thresholds at 80% correct choices for the four conditions. We also calculated the differences between the contrast thresholds of the pesticide groups and between the contrast thresholds of the motivation groups. All these calculations were bootstrapped five hundred times to obtain their 95% confidence intervals.

To analyse the latency to make the first choice, we first log_10_ transformed the latencies to limit the effect of extreme values. We then fitted a model with the transformed latencies as the dependent variable and a Student-t family (t_family in glmmTMB) and identity link function. Here again, we included target irradiance, motivation group, and pesticide group and their interactions as independent variables and bee identity as a random effect. We also modelled the dispersion of the latencies depending on the log transformed target irradiance, the pesticide group and the motivation group, using the dispersion formula (dispformula) in glmmTMB. Due to recording issues, 5 tests in the non-pesticide high motivation group and 3 in the low motivation group could not be included in our analysis.

We used likelihood ratio tests with the anova() function in R to determine which dependent variable had a significant effect on the dependent variable. We started from the full model (with all variables and interactions) and removed one by one each non-significant interaction/variable, comparing each new model with the previous one with this interaction/variable (See Table 1 for model comparisons).

We then examined the estimates of the model that included only the significant variables and interactions to understand their effect on the dependent variable (Full results in Table 2).

## Results

### Choice accuracy

When analysing the first six choices, model comparisons showed that the best model included main effects of the motivation group, the pesticide treatment and the target irradiance but not their interactions (Table 1A). Bees made significantly more accurate choices when the targets became more visible (with higher target irradiance) (Fig. 2A ; Table 2A ; Estimate ± Standard Error = 0.956 ± 0.065, Z = 14.750, p < 0.001). Bees from the low motivation group were also significantly more likely to choose locations without a target than bees from the high motivation group (Estimate ± S.E. = - 0.303 ± 0.136, Z = -2.220, p = 0.026). This could indicate that the lower motivation group had a lower attentional state than the other group leading to worse target detection. Interestingly, the imidacloprid treatment also significantly lowered the bees’ accuracy in their six first choices across all target irradiances and for both motivation groups (Estimate ± S.E. = -0.382 ± 0.144, Z = -2.663, p = 0.008).

**Table 1:** Model comparisons analysing the influence of variables and interactions on bee choice accuracy (probability to choose target locations) in A) their first six choices and B) their first choice, and C) latency to make their first choice. Model comparisons analysing the influence of variables and interactions on the latency to make this first choice. Each line shows the result of the test comparing a model including the tested variable with a model without it. The first line of each sub-table (A-C) used the full model as baseline against which the model without the 3-way interaction was compared. Each non-significant variable was excluded from all subsequent compared models. The significance of each effect in the first column was then tested by removing it from the model and comparing the models. The table shows the results, degree of freedom and p-values of the likelihood ratio tests used for the analyses. Significance levels are as follows: * <0.05, **<0.01, ***<0.001.

| A) | Analysis of choice accuracy in the first six choices |  |  |  |
| --- | --- | --- | --- | --- |
| Tested effect | $\chi^2$ | df | P | Significance |
| Log(Target irradiance) : Motivation group : Pesticide group | 0.061 | 1 | 0.805 |  |
| Motivation group : Pesticide group | 0.146 | 1 | 0.703 |  |
| Log(Target irradiance) : Pesticide group | 0.541 | 1 | 0.462 |  |
| Log(Target irradiance) : Motivation group | 3.115 | 1 | 0.078 |  |
| Motivation group | 4.815 | 1 | 0.028 | * |
| Pesticide group | 6.579 | 1 | 0.010 | * |
| Log(Target irradiance) | 239.53 | 1 | < 0.001 | *** |
| B) | Analysis of the first choice accuracy |  |  |  |
| Tested effect | $\chi^2$ | df | P | Significance |
| Log(Target irradiance) : Motivation group : Pesticide group | 1.365 | 1 | 0.243 |  |
| Motivation group : Pesticide group | 1.553 | 1 | 0.213 |  |
| Log(Target irradiance) : Motivation group | 0.154 | 1 | 0.695 |  |
| Log(Target irradiance) : Pesticide group | 2.925 | 1 | 0.087 |  |
| Motivation group | 1.922 | 1 | 0.166 |  |
| Pesticide group | 3.085 | 1 | 0.079 |  |
| Log(Target irradiance) | 62.521 | 1 | < 0.001 | *** |
| C) | Analysis of the latency to first choice (s) |  |  |  |
| Tested effect | $\chi^2$ | df | P | Significance |
| Log(Target irradiance) : Motivation group : Pesticide group | 0.598 | 1 | 0.439 |  |
| Motivation group : Pesticide group | 1.930 | 1 | 0.165 |  |
| Log(Target irradiance) : Motivation group | 2.630 | 1 | 0.105 |  |
| Log(Target irradiance) : Pesticide group | 3.713 | 1 | 0.054 |  |
| Pesticide group | 2.982 | 1 | 0.084 |  |
| Motivation group | 4.207 | 1 | 0.040 | * |
| Log(Target irradiance) | 139.690 | 1 | < 0.001 | *** |

We used the values predicted by the best (Table 2A) model to calculate the contrast sensitivity thresholds of the bees for 80% accuracy in the first six choices. We found that in the no pesticide conditions, bees achieved 80% accuracy with targets at 0.0186 µW/cm^2^/nm (95% C.I. [0.0145, 0.0231]) target irradiance in the high motivation group and 0.0255 µW/cm^2^/nm (95% C.I. [0.0203, 0.0313]) in the low motivation group (Fig. 3). This represented a significant -0.007 µW/cm^2^/nm (95% C.I. [-0.0140, -0.0012]) difference of contrast sensitivity. In the pesticide conditions, the high motivation group reached 80% accuracy at 0.0278 µW/cm^2^/nm (95% C.I. [0.0200, 0.0373]) target irradiance while the low motivation group reached the same accuracy at 0.0381 µW/cm^2^/nm (95% C.I. [0.0279, 0.0489]) (Fig. 4). This -0.0103 µW/cm^2^/nm (95% C.I. [-0.0211, -0.0017]) difference in contrast sensitivity was also significant. When comparing the pesticide and non-pesticide group, we found a significant -0.0091 µW/cm^2^/nm (95% C.I. [-0.0184, -0.0019]) decrease in contrast sensitivity in the high motivation bees. Similarly, there was a significant -0.0126 µW/cm^2^/nm (95% C.I. [-0.0248, -0.0025]) decrease in the low motivation group.

We also analysed the accuracy of just the first choice of the bee in each test trial. This is likely the choice during which the bees made their choice the furthest from the screen and thus at the point when target detection was the most difficult. Model selection showed that the best model only included the target irradiance as an independent variable (Table 1B). This model showed that, as for when examining all first six choices, choice accuracy in the first choice increased with the irradiance of the targets (Fig. 2B ; Table 2B ; Estimate ± S.E. = 1.470 ± 0.217, Z = 6.790, p < 0.001).

**Table 2:** Results of the selected models for A) choice accuracy in the first six choices, B) choice accuracy of the first choice and C) latency to make the first choice. The table shows the estimates, standard errors, Wald test results and p-values for each variable and interaction in the models. Significance levels are: * <0.05, **<0.01, ***<0.001.

| A) Model |  | Choices accuracy ~ Log(Target irradiance) * Motivation group + Pesticide group + (1 \| Bee ID) |  |  |  |
| --- | --- | --- | --- | --- | --- |
| Variable | Estimate | Standard Error | Z | P | Significance |
| Intercept | 5.194 | 0.254 | 20.487 | < 0.001 | *** |
| Log(Target irradiance) | 0.956 | 0.065 | 14.750 | <0.001 | *** |
| Motivation group (low motivation) | -0.303 | 0.136 | -2.220 | 0.026 | * |
| Pesticide group (Imidacloprid) | -0.382 | 0.144 | -2.663 | 0.008 | ** |
| B) Model |  | Choice accuracy ~ Log(Target irradiance) + (1 \| Bee ID) |  |  |  |
| Variable | Estimate | Standard Error | Z | P | Significance |
| Intercept | 7.267 | 0.861 | 8.438 | <0.001 | *** |
| Log(Target irradiance) | 1.470 | 0.217 | 6.790 | < 0.001 | *** |
| C) Model |  | Log10(Latency) ~ Log(Target irradiance) + Motivation group + (1 \| Bee ID) |  |  |  |
| Variable | Estimate | Standard Error | Z | P | Significance |
| Intercept | 0.287 | 0.037 | 7.760 | < 0.001 | *** |
| Log(Target irradiance) | -0.166 | 0.013 | -12.902 | <0.001 | *** |
| Motivation group (low motivation) | 0.053 | 0.025 | 2.091 | 0.037 | * |

Contrary to what was seen for first choice accuracy, the motivation group influenced first choice latency (Table 1C). Target irradiance also had an effect as expected but not pesticide treatment. Here again, none of the variables interacted with each other. The best model of our data showed that bees took less time to make their first choice as the irradiance of the targets increased and they became more visible (Fig. 2C ; Table 2C ; log10(Estimate ± S.E). = -0.166 ± 0.013, Z = -12.902, p < 0.001). First choice latency was also longer for the low motivation group than the high motivation group (log10(Estimate ± S.E). = 0.053± 0.025, Z = 2.091, p = 0.037). In experiments testing for attention, reaction time is often a key measure of attention (Bichot and Schall, 2002; Jigo and Carrasco, 2018; Mahadevan et al., 2018; Pham et al., 2018; Posner, 1980; Quest et al., 2022; Sridharan et al., 2014; Wegener et al., 2008). Our results could therefore indicate again, that the high motivation group had increased attention during their search for the targets, leading to increased perceived target contrast and detection.

**Figure 2:**
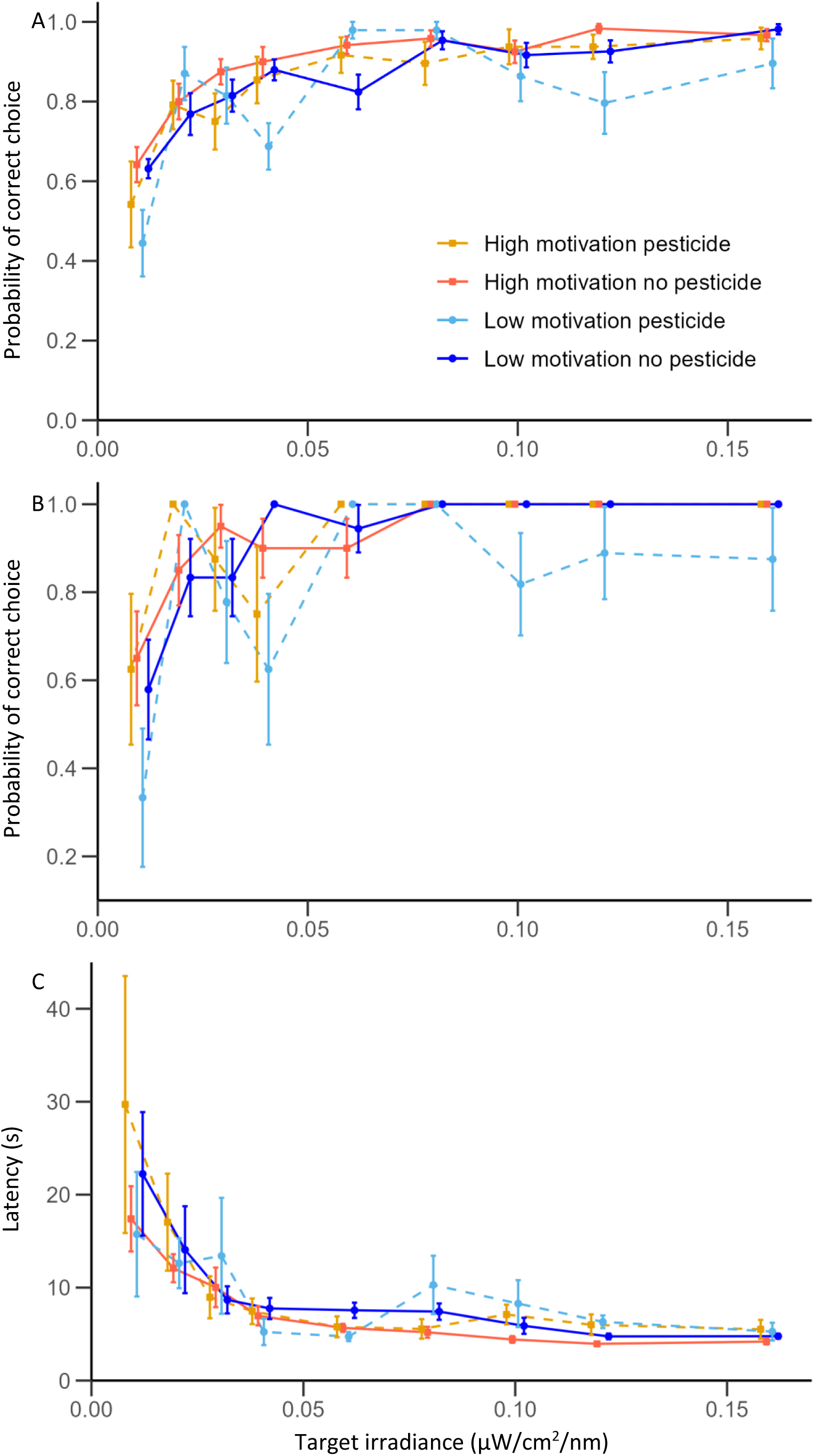
Effects of motivation, pesticide and target irradiance on A) bee choice accuracy (probability to choose a target location) in their first six choices B) choice accuracy in their first choice and C) their latency to make the first choice. The points are average values and error bars show the standard errors. The data for the high motivation group is presented in orange for the experiment without pesticide and in yellow for the experiment with pesticide. The data of the low motivation group is shown in dark blue for the experiment without pesticide and light blue for the experiment with pesticide. Data from the pesticide experiment are shown with dashed lines.

**Figure 3:**
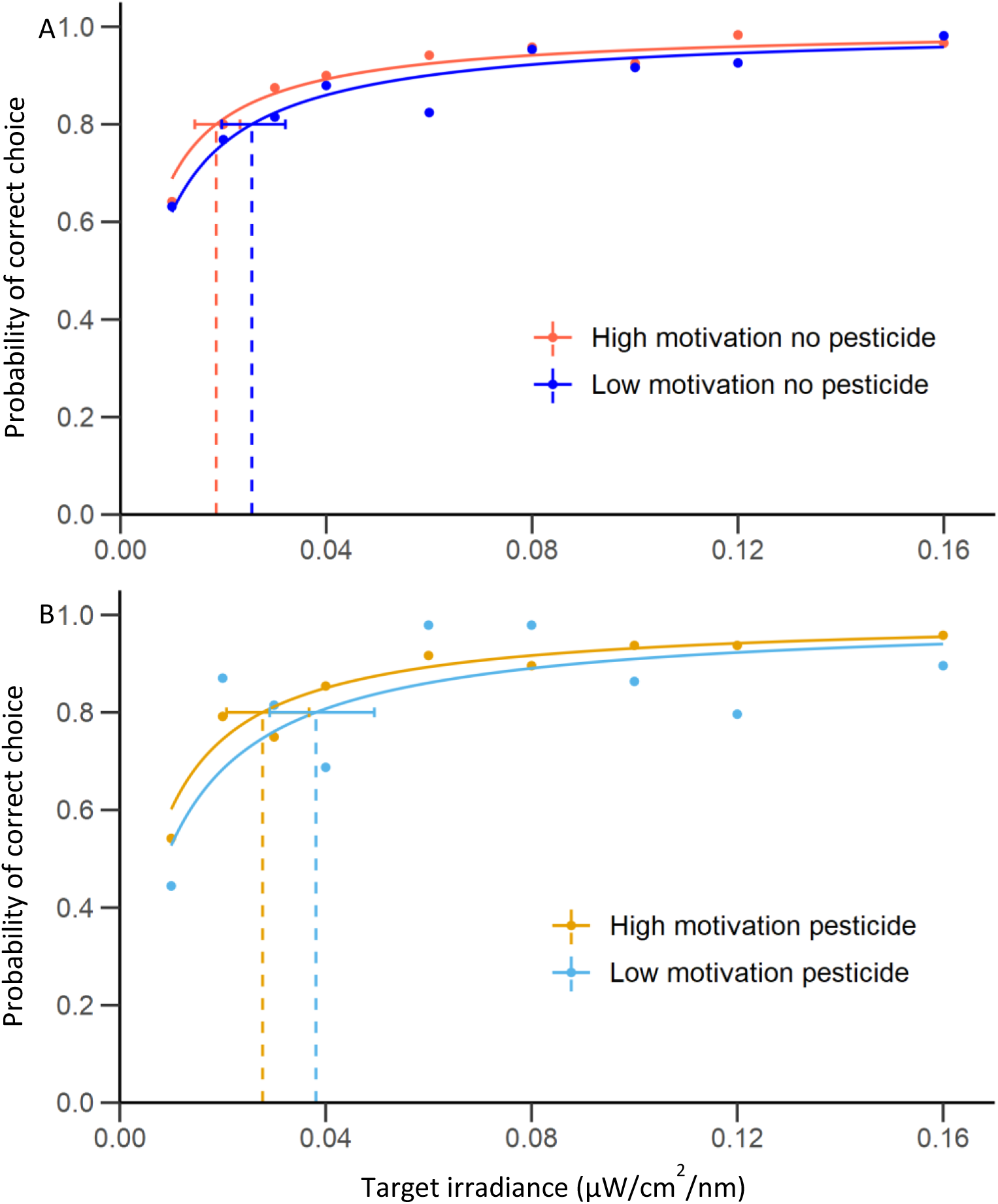
Comparisons of the contrast sensitivity of the high and low motivation groups in A) the no pesticide and B) the pesticide condition. The curves are the contrast sensitivity curves predicted by the best model fitting the first six choices accuracy data (Table 2A). The dots represent the average probability of correct choices. Dashed lines show the target contrast threshold of each group to reach 80% accuracy. The error bars show the 95% confidence intervals of these thresholds. The colour code is similar to figure 2.

**Figure 4:**
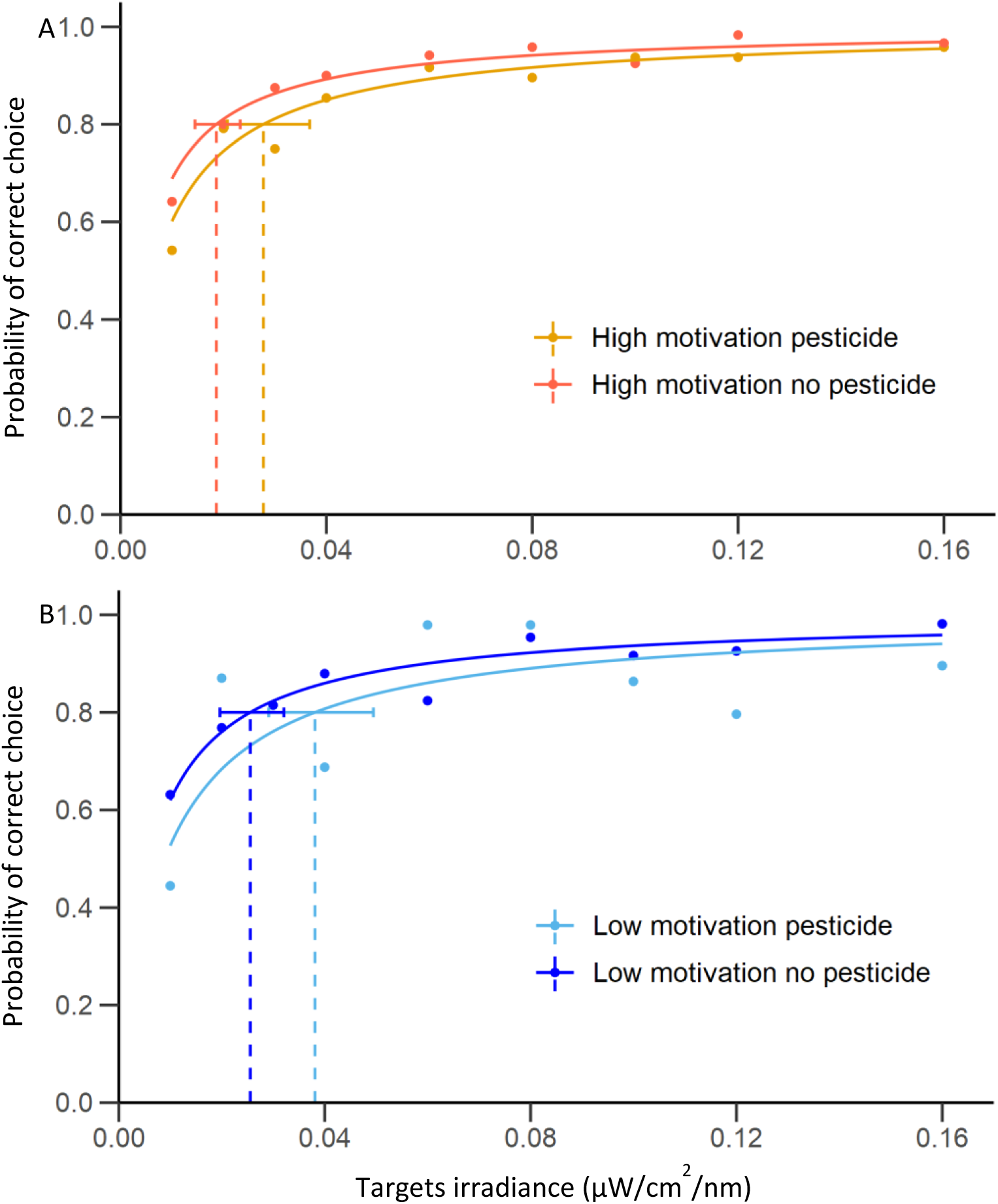
Comparisons of the contrast sensitivity of the pesticide and no pesticide groups in A) the high motivation and B) low motivation bees. The curves are the contrast sensitivities predicted by the model best fitting the first six choice accuracy data (Table 2A). The dots are the mean probability of correct choices. The dashed lines are the contrast threshold of the bees for 80% accuracy and the error bars show their 95% confidence intervals. The colours are the same than in figure 2.

## Discussion

Our results showed that bees were more likely to accurately choose targets with increases in target contrast, as expected. This is consistent with previous studies that have shown the importance of the contrast of a target or a pattern for the ability of bee to detect them (Chakravarthi et al., 2016; Procenko and Nityananda, 2026; Robert et al., 2026). This therefore allowed us to ask more detailed question about the factors influencing contrast sensitivity.

### The effect of reward

Our reward treatment influenced both the probability that our bees made accurate choices in their first six choices and their latency to make their first choice in each trial. Choice accuracy and reaction time are classic measures of attention in experiments with humans and non-human primates. Our finding thus suggests that higher reward induced increased motivation and attention, leading to increased contrast sensitivity and enhanced detection of the targets. This is comparable to what has been reported in humans. Reward quality increases attention toward the characteristics of a visual stimulus associated with this reward or the location where it could appear (Anderson et al., 2011; Bucker and Theeuwes, 2017; Della Libera and Chelazzi, 2009; Libera and Chelazzi, 2006; Massar et al., 2016; Raymond and O’Brien, 2009). Feature-based attention has also been shown to increase contrast sensitivity (Herrmann et al., 2012).

Surprisingly, we did not find an effect of our motivation group on the first choice accuracy of the bees. One explanation for this could be a lack of power in our analysis of the first choice as this variable had fewer datapoints. In addition, our original assumption was that the effect of attention would be more measurable during the first choice of each trial because it was possibly the most difficult one for the bees as they presumably made it further away from the screen. However, it is also possible that the first choice may not have been as difficult as we previously thought. Indeed, when making this choice, bees still had six correct locations available to visit. Assuming that the insects avoided revisiting the same location, finding the remaining targets may have become increasingly difficult after each correct choice. In this case, finding the last couple of targets may have been more difficult than making a correct first choice.

In addition, even if the first choice was really the most difficult one to make for the bees, our animals were flying freely and could approach the screen without any time constraint. Therefore, the effect of attention may not be seen on the first choice accuracy but on the latency to make this first choice. If low motivation bees had more trouble detecting the targets than the high motivation ones due to lower contrast sensitivity, they could take more time to make their first choice. This is, in fact, what we observed in our experiment. The latency to make this first choice was shorter for the high motivation group than for the low motivation one. This difference could support the idea that the high motivation bees had higher contrast sensitivity, leading them to need less time to detect one of the targets. This may be compared to the shorter reaction times of human subjects and other vertebrates during target detection tasks after being cued to the correct area of their visual field or after learning a feature of the target (Bichot and Schall, 2002; Jigo and Carrasco, 2018; Mahadevan et al., 2018; Pham et al., 2018; Quest et al., 2022; Sridharan et al., 2014; Wegener et al., 2008).

Our results add to the growing neurological and behavioural evidence that insects possess cognitive processes similar to vertebrate attention (Lancer et al., 2019; Lancer et al., 2022; Nityananda, 2016; Robert et al., 2024; Robert et al., 2025; Sareen et al., 2011). The majority of these studies in insects demonstrated the existence of a bottom-up form of attention. This was shown in predatory insects for which rapidly detecting the sudden movements of a prey could improve their capture success (Lancer et al., 2019; Lancer et al., 2022; Robert et al., 2025), or in defenceless prey such as fruit flies (Sareen et al., 2011) where quickly reacting to the sudden approach of a predator could increase their chances of survival. To our knowledge, the only work attempting to find bottom-up attention in bees using a classic cuing paradigm was unsuccessful (Robert et al., 2026). However, some studies have indicated that bees may have a process comparable to top-down attention. For example, after having associated a type a flower with a high reward, bumblebees increase the proportion of time spent visually scanning these flowers (Nityananda and Chittka, 2021; Robert et al., 2024). Our current study would support this conclusion.

There could be some other potential explanations for our results. The difference in choice accuracy between our motivation group could reflect a difference in foraging persistence when bees found no reward during the tests. Bees tend to switch to different flowers if the quality of the reward received decreases or the reward disappears altogether (Hemingway and Muth, 2022; Townsend-Mehler and Dyer, 2012; Townsend-Mehler et al., 2011; Waldron et al., 2005; Wiegmann et al., 2003). Honeybees are also more likely to persist on flowers that they perceived as providing high quantity of nectar (Gil et al., 2007). Another study have shown that, during unrewarded tests, bees were less likely to switch from visiting flowers they had associated with 50% sucrose rewards if the alternative flowers were flowers associated with 30% sucrose, than if the alternative flowers were also associated with 50% sucrose (Nityananda and Chittka, 2021). Therefore, if bees estimated the reward differential between flower types to make such decision to switch, it is possible that bees trained with 50% sucrose were giving up looking for the targets later than the bees trained with 30% sucrose.

A difference in foraging persistence due to a difference in motivation would also not explain why the high motivation group made their first choice faster than the low motivation one. Studies have shown that bees foraging for food sources with high sugar concentration fly faster than bees foraging for food sources with lower sugar concentration (Baciadonna and Nityananda, 2026; Procenko et al., 2024; V. Frisch and Lindauer, 1955; Willemet, 2024) presumably due to a higher motivation. This could explain the differences in first choice latency between motivational groups. However, is it also possible that this higher flight speed is enabled by a higher visual sensitivity due to an increase in the bees’ attentional state. This would be comparable to faster reaction times in experiments with humans and non-human primates (Bichot and Schall, 2002; Jigo and Carrasco, 2018; Mahadevan et al., 2018; Pham et al., 2018; Posner, 1980). This is a promising possibility, and future studies could confirm this using neurobiological techniques.

### The effect of pesticide

The effect of imidacloprid on the choice accuracy could be explained by either an effect on motivation and attention, or potentially an effect on the visual learning and memory of the bees. A few studies have reported that sublethal doses of imidacloprid disturbed visual processes in locusts or wasps (Corcoran and Tibbetts, 2023; Parkinson and Gray, 2019; Parkinson et al., 2017).

Imidacloprid also appears to affect olfactory learning in bees but may not affect their visual learning (Aguiar et al., 2023; Decourtye et al., 2003; Decourtye et al., 2004a; Decourtye et al., 2004b; Iqbal et al., 2019; Li et al., 2019; Mengoni Goñalons and Farina, 2015; Zhang and Nieh, 2015; for a review on how neonicotinoids affect bee olfactory learning, see Paoli and Giurfa, 2024). To our knowledge, only one study showed that imidacloprid hindered visual learning in bumblebees (Paus-Knudsen et al., 2023), but in this experiment, bees were exposed to the pesticide for nine days. In our experiment, there is some possibility that the bees may also have been exposed to the pesticide for several days. Indeed, we cannot exclude that they may have consumed the contaminated sugar solution collected and brought back to the colony by previously tested bees. However, the amount of contaminated sugar solution collected by the few bees that participated in our pesticide experiment is certainly negligible compared to the amount of pesticide free sugar solution collected by the entire colony during evenings and weekends. Therefore, it seems unlikely that this little amount of pesticide had a significant enough impact on our tested bees to affect their visual learning and memory. The effect of imidacloprid on our tested bees was more likely induced by the direct consumption of the sugar rewards during our training and refresher bouts. This direct exposure was much shorter as most bees completed the full experiment within a single day with only one bee finishing her last test 28 hours after her first training.

If individual bees’ exposure to imidacloprid in our experiment was indeed short, it may not have been enough to directly affect bee visual learning. In their study Muth and colleagues trained bumblebees to collect a reward on artificial flowers of a specific colour and odour and to ignore flowers of different colour and odour (Muth et al., 2019). The tests presented the same two flower types on which the bees were trained. They also presented two flower types where the flower either had the correct colour but wrong odour or vice versa. The results of this experiment showed that bees dosed with a single sublethal dose of imidacloprid before the training phase differed from control bees only in their number of choices to forage on flowers with the right colour but the wrong odour. This result indicated that the pesticide treatment may have disturbed either the bumblebees scent learning or olfaction in general but not their visual learning. Additional studies showed that a short exposure to imidacloprid does not affect visual learning in bees but reduces the motivation to forage (Karahan et al., 2015; Muth and Leonard, 2019). Therefore, it is likely that our imidacloprid treatment had the same effect as the difference in rewards, in that it affected foraging motivation. Decreasing motivation could have decreased attention and thus the ability to detect the targets. This would explain why imidacloprid decreased choice accuracy in our experiments.

Despite the decrease in accuracy, we do not see a corresponding decrease in first choice latency in our pesticide treated bees. The most plausible explanation is, here again, the fact that our sample sizes were quite small. As can be seen in Figure 2C, there is a lot of variation in the data from the pesticide groups which suggests that our sample size was not large enough to detect any effects induced by imidacloprid. Alternatively, imidacloprid may have affected bumblebee attention in a different way than our motivation treatment.

Imidacloprid disturbs the cholinergic neural pathways of insects by binding to nicotinic acetylcholine receptors (Brown et al., 2006; Matsuda et al., 2020; Taillebois et al., 2018). In mammals, numerous studies have shown that cholinergic circuits are key to the deployment of top-down attention (for reviews see Hasselmo and Sarter, 2011; Klinkenberg et al., 2011; Parikh and Bangasser, 2020; Villano et al., 2017). If bumblebee attention functions in a similar way, it is possible that the imidacloprid treatment directly affected the bees’ ability to tune their sensory system when searching for the targets, by disturbing their cholinergic neural circuits. Cholinergic circuits in the prefrontal cortex of rats have been shown to be critical to sustained attention during demanding attentional tasks (Dalley et al., 2004; Newman and McGaughy, 2008). In an experiment, rats with lesioned cholinergic neurones in their prefrontal cortex were trained to perform a visual detection task. During tests, auditory distractor sounds were played. If the distractor sound was rhythmical, it did not affect the animals’ ability to detect the visual target. However, an arguably more demanding attentional condition with an irregular (unpredictable) auditory distractor significantly reduced visual target detection (Newman and McGaughy, 2008). In our experiments, it is possible that the first choice was an easier task compared to subsequent choices, when bees had to maintain their attention to detect the remaining targets and simultaneously filter out the locations they had already visited. This might explain why we don’t see the effect of imidacloprid on first choice latency but found one on the accuracy in the first six choices.

Overall, our study is a first step in demonstrating that bees possess a cognitive process similar to vertebrate top-down attention and that this process may increase their contrast sensitivity. It also suggests that imidacloprid may have an effect on this process, presumably through its impact on bee motivation or the cholinergic receptors that may underlie attention-like processes. However, more work will be needed to confirm our findings and disentangle the effect of motivation on this process from the other effects it can have on bee behaviour, such as their persistence to forage. Neurological studies may allow such disambiguation. In the brain of primates, it is possible to record the tuning of the sensory system in order to better detect the features of the target the animal is looking for (David et al., 2008; Hayden and Gallant, 2005; Treue and Trujillo, 1999; Zhou and Desimone, 2011; for reviews, see Liu, 2019; Maunsell and Treue, 2006). In dragonflies, studies have been able to record a neuron selectively reacting to the movement of a target and ignoring other targets if the one selected was primed before starting moving (Lancer et al., 2019; Lancer et al., 2022). We therefore recommend that future studies investigate how bees may tune their sensory system to better detect the features associated with profitable food sources.

## Data Availability

Data and statistical analyses code are made available at: https://figshare.com/s/93cafd99d774b99b115f

## Author Contributions

V.N. obtained the funding. V.N. and T.R. designed the experiments. E.F, H.L and T.R. conducted the experiments. H.L. analysed the videos. T.R. ran the statistical analyses and wrote the paper. V.N. edited the manuscript. M.N. conducted a pilot experiment.

## Acknowledgment

For the purpose of open access, the authors have applied a Creative Commons Attribution (CC BY) licence to any Author Accepted Manuscript version arising.

